# A compartment size dependent selective threshold limits mutation accumulation in hierarchical tissues

**DOI:** 10.1101/719575

**Authors:** Dániel Grajzel, Imre Derényi, Gergely J. Szöllősi

**Affiliations:** MTA-ELTE “Lendület” Evolutionary Genomics Research Group, Pázmány P. stny. 1A, H-1117 Budapest, Hungary; Department of Biological Physics, Eötvös Loránd University, Pázmány P. stny. 1A, H-1117 Budapest, Hungary; MTA-ELTE Statistical and Biological Physics Research Group, Hungarian Academy of Sciences, H-1117, Budapest, Hungary; Evolutionary Systems Research Group, Centre for Ecological Research, Hungarian Academy of Sciences, 8237 Tihany, Hungary

**Keywords:** Somatic Evolution, Tissue hierarchies, Cancer Evolution, Physics of Cancer

## Abstract

Cancer is a genetic disease fueled by somatic evolution. Hierarchical tissue organization can slow somatic evolution by two qualitatively different mechanisms: by cell differentiation along the hierarchy “washing out” harmful mutations (Nowak et al. 2003, Werner et al. 2013) and by limiting the number of cell divisions required to maintain a tissue (Derényi and Szöllő si 2017). Here we explore the effects of compartment size on somatic evolution in hierarchical tissues by considering cell number regulation that acts on cell division rates such that the number of cells in the tissue has the tendency to return to its desired homeostatic value. Introducing mutants with a proliferative advantage we demonstrate the existence of a third fundamental mechanism by which hierarchically organized tissues are able to slow down somatic evolution. We show that tissue size regulation leads to the emergence of a threshold proliferative advantage, below which mutants cannot persist. We find that the most significant determinant of the threshold selective advantage is compartment size, with the threshold being higher the smaller the compartment. Our results demonstrate that in sufficiently small compartments even mutations that confer substantial proliferative advantage cannot persist, but are expelled from the tissue by differentiation along the hierarchy. The resulting selective barrier can significantly slow down somatic evolution and reduce the risk of cancer by limiting the accumulation of mutations that increase the proliferation of cells.

Significance Statement
Renewed tissues of multicellular organism accumulate mutations that lead to ageing and cancer. To mitigate these effects self-renewing tissues produce cells along differentiation hierarchies, which have been shown to suppress somatic evolution both by limiting the number of cell divisions, and thus reducing mutational load, and by differentiation “washing out” mutations. Our analytical results reveal the existence of a third mechanism: a compartment size dependent threshold in proliferative advantage, below which mutations cannot persist, but are rapidly expelled from the tissue by differentiation. In sufficiently small compartments the resulting selective barrier can greatly slow down somatic evolution and reduce the risk of cancer by preventing the accumulation of mutations even if even they confer substantial proliferative advantage.

---

**T**umors develop as genetic and epigenetic alterations spread through a population of premalignant cells and some cells accumulate changes over time that enable them and their descendants to persist within tissues (1, 2). From an evolutionary perspective each tumor is an independent realization of a common reproducible evolutionary process involving “adaptive” mutations that are preferentially selected by the tumor environment. This process is clonal, which means that a subset of mutations termed “drivers” confer clonal growth advantage and they are causally implicated in cancer development.

A large body of work (2–5) has focused on understanding clonal evolution of an initially homogeneous populations of identical cells, a subset of which progress toward cancer as they accrue driver mutations. Beerenwinkel et al. (6), for instance, considered the Wright-Fisher process (a homogeneous population of initially identical cells) to explore the basic parameters of this evolutionary process and derive an analytical approximation for the expected waiting time to the cancer phenotype and highlighted the relative importance of selection over both the size of the cell population at risk and the mutation rate.

Self-renewing tissues, which must generate a large number of cells during an individual’s lifetime and in which tumors typically arise, are comprised of a hierarchy of progressively differentiated cells and, as a result, are not homogeneous populations of identical cells. There is empirical evidence (7–9) and theoretical rationale (10–12) that such hierarchical tissue architecture has profound effect on neoplastic progression. Theoretical work has demonstrated that hierarchically organized tissues suppress tumor evolution by limiting the accumulation of somatic mutations in two fundamentally different ways, as follows:

As described in a seminal paper by Nowak et al. (11) the linear flow from stem cells to differentiated cells to apoptosis in a spatially explicit, strictly linear organization has the property of canceling out selective differences. Nowak et al. considered a system, where only asymmetric cell divisions are allowed, i.e., after each cell division one of the daughter cells differentiates to the next level of the hierarchy pushing all cells at higher levels further along the hierarchy (see Fig. 1a). In this idealized construction mutations, irrespective of how much they increase division rate, are invariably “washed out” unless they occur in the stem cell at the root of the hierarchy. In a more general setting, where symmetric divisions are allowed, the strength of this “washing out” effect can be quantified by introducing the self-renewal potential of cells. The self-renewal potential is defined as the logarithm of the ratio between the rate of cell divisions that increase the number of cells at a given level of the hierarchy (division producing two cells at the same level) and the rate of events that result in the reduction at the level (division producing two differentiated cells that move higher up in the hierarchy or cell death). h1 healthy homeostatic tissues the self-renewal potential of stem cells is zero (corresponding to equal rates of differentiation and self-renewal), while for differentiated cells it is always negative, as these cells have an inherent proliferative disadvantage as a result of which they are eventually “washed out” of the tissue from cells differentiating from lower levels of the hierarchy. In the following, lower (higher) refers to levels closer to (further away from) the stem cell compartment.

**Fig. 1.**
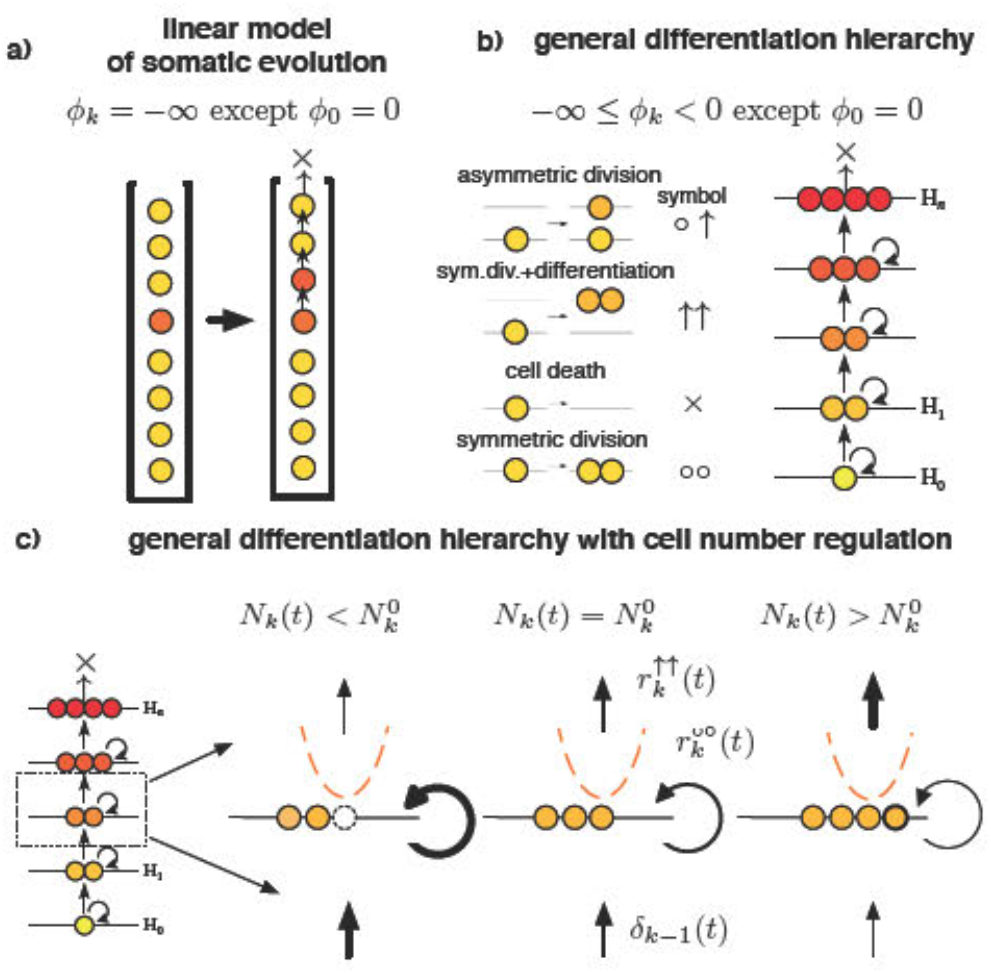
Self-renewing tissue comprised of a hierarchy of progressively differentiated cells can suppress somatic evolution. a) The linear process of somatic evolution considers a strict linear organization, where after each cell division one of the daughter cells d fferentiates to the next level pushing all cells above further along and the top most cell is lost from the system. Such an idealized construction, where self.renewal at individual levels of the hierarchy is not allowed has a minimal self-renewal potential *ϕ*_*k*_ = − ∞, with the exception of the stem cell level at the root of the hierarchy with *ϕ*_0_ = 0. This has the effect of canceling out selective differences between cells, i.e., any non stem cell, regardless of how large its division rate is, will be “washed out” of the tissue by cell differentiating from below. b) General differentiation hierarchies are characterized by intermediate values of the self-renewal potential with the exception of the stem cell. In such systems, in the absence of cell number regulation, any mutant with a proliferative advantage, i.e., a positive self-renewal potential, will spread exponentially if it dces not go extinct stochastically. c) We introduce cell number regulation that changes the rate of different events such that the strength and direction of the regulation depends on the difference between the number of cells present at a given time *N*_*k*_(*t*) and the homeostatic number 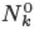 in a manner equivalent to being in a quadratic potential (cf. Eq. (3)). As described in the text, this leads to the emergence of a positive threshold proliferative advantage below which mutants cannot persist.

More recently, Derényi and Szöllősi (12) showed that in self-renewing tissues hierarchical organization provides a robust and nearly ideal mechanism to limit the divisional load (the nwnber of divisions along cell lineages) of tissues and, as a result, minimize the accumulation of somatic mutations. The theoretical minirnwn nwnber of cell divisions can be very closely approached: as long as a sufficient number of progressively slower dividing cell types towards the root of the hierarchy are present, optimal self-sustaining differentiation hierarchies can produce *N* terminally differentiated cells during the course of an organism’s lifetime from a single precursor with no more than log_2_(*N*) + 2 cell divisions along any lineage.

Here, we examine the effect of compartment size by introducing interaction among cells in the form of cell number regulation, which acts on the cell division rates such that the nwnber of cells at each hierarchical level of the tissue has the tendency to return to its desired homeostatic value. We consider a single (non stern cell) level of the hierarchy that is renewed from below by cell differentiation. We introduce mutants with a proliferative advantage, i.e., mutants with a positive self-renewal potential. As detailed below, using both simulations and an approximation adopted from nonequilibrium statistical physics, we find that under a wide range of parameters a third fundamental mechanism exists by which hierarchically organized tissues can slow down somatic evolution and delay the onset of cancer.

## 1. Results

We consider level ***k*** > 0 of a general differentiation hierarchy that is renewed by cell differentiation from level ***k*** − 1 below. The tissue dynamics is described by the rates of asymmetric differentiation (ο↑), symmetric division with differentiation (↑↑), symmetric division (οο), and cell death (×) (see Fig. 1 b and c).

At homeostasis (i.e., when the number of cells, ***N_k_***, at each level coincides with its homeostatic value, 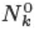) the evolutionary dynamics of level ***k*** is determined by the per cell rate 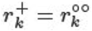 of cell nwnber increase through symmetric cell division (οο), the per cell rater 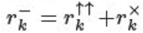 of cell number decrease through either symmetric division with differentiation (↑↑) or cell death (×), and the per level rate 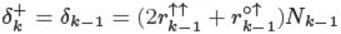 of cell number increase through differentiation from below. In the following, we focus on a single (non stem cell) level, and drop index ***k*** for brevity. Homeostasis implies that the rates satisfy the stationarity condition

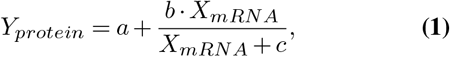

To model tissue size regulation we consider a generic cell number-dependent regulation scheme that acts to return the number of cells in the compartment to its homeostatic value. Biologically such a regulation scheme corresponds to, e.g., the local concentration of a regulatory signal that conveys information on the density of cells in a compartment. To formalize cell-number regulation we introduce cell number-dependent multiplicative rate modifiers *ρ*^+^ (*N*) and *ρ*^−^(*N*) for, respectively, events that increase and decrease cell number. Maintaining homeostasis requires that *ρ*^+^ (*N*) > 1 and *ρ*^−^(*N*) < 1 if *N* < *N*^0^; and *ρ*^+^(*N*) < 1 and *ρ*^−^(*N*) > 1 if ***N*** > *N*^0^. These rate modifiers define an abstract confining potential ***U(N)*** up to an additive constant:

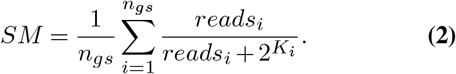

with a mini:mwn at ***N*** = ***N***^0^. For mathematical convenience, we approximate the confining potential with a parabolic (harmonic) form:

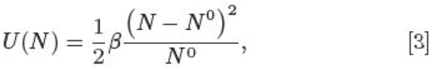

characterized by a single parameter (potential strength) *β*.

The role of the confining potential is to limit the variance of the number of cells to ***N^0^***/*β*. It is this confining potential that plays the most significant role in slowing down somatic evolution, as shown below.

The particular choice of how the confining potential is distributed between the cell number increasing and decreasing rate modifiers (to satisfy Eq. (2)) has only marginal effect on the dynamics. Here, for simplicity, we make the symmetric choice:

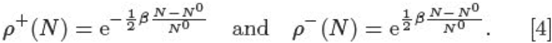

We measure time in units of 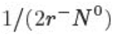. Thus, the life-time of the tissue ***T*** is identical to the expected number of differentiated cells that would be produced by this level under homeostatic conditions during the individual’s lifetime, if all the cell number decreasing events were symmetric cell differentiations. As the asymmetric differentiations (ο↑) do not have any influence on the number of cells of this level, we set its rate (*r*^ο↑^) to zero for convenience.

The self-renewal potential of the cells in a healthy homeostatic tissue is defined as

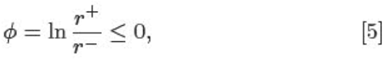

which converges to −∞ as the rate of the (οο) events approaches zero.

We introduce mutants with an elevated rate of divisions that increase cell number, 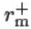, such that it exceeds the rate of cell number decrease: 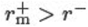. This corresponds to a positive self-renewal potential for mutant cells:

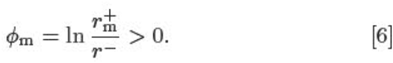

In the absence of cell nun1ber regulation (i.e., *β* = 0) a single such mutant either goes extinct stochastically, or spreads exponentially with probability (13)

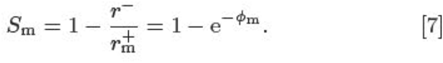

In the following we use *S*_*m*_ to parametrize the selective advantage of mutants. We note that in the absence of differentiation from below (i.e., ***δ***^+^ = 0), for fixed population size (i.e., *β* ➔ ∞), for all but extremely small populations or nearly neutral mutations, *S*_*m*_ also corresponds to the probability of fixation of the mutant (14–16).

Denoting the number of mutant cells by ***N_m_*** and wild type cells by ***N_w_*** the dynamics is described by the transition rates

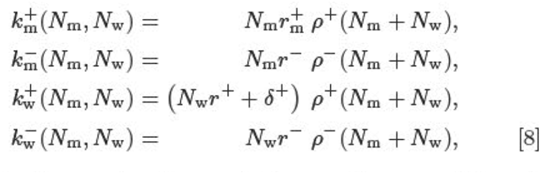

where the lower index (m or w) denotes the type of the cell (mutant or wild, respectively), and the upper index indicates whether the number of cells of the given type is increased (+) or decreased (−). The transition rates can be shown to correspond to a reversible Markov process in the effective potential

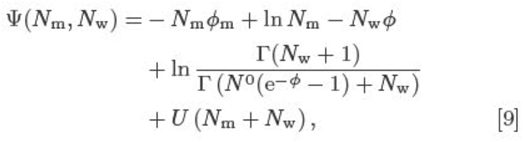

where Г represents the gamma function.

The continuous interpolation of this potential is shown in the bottom panels of Fig. 2 a and b for different parameters. An effective potential, such as Eq. (9), can always be defined if the mutant and wild type transition rates depend only on the number of cells of the given type, and cell number regulation - which acts as a multiplicative modifier of these rates - depends only on the total number of cells (see Supplementary Appendix).

**Fig. 2.**
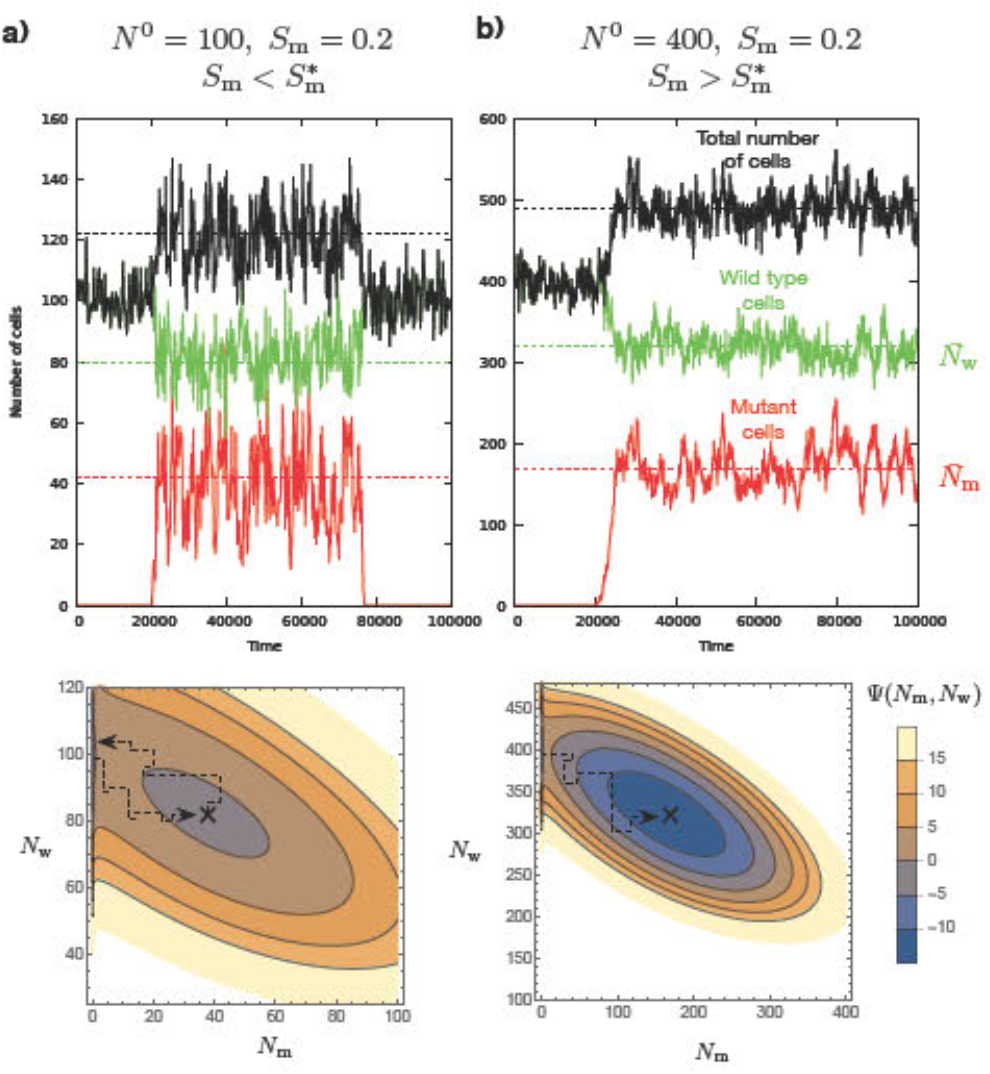
Mutants go extinct under a threshold proliferation advantage. The continuous lines show the size of the mutant (red) and wild type (green) populations and their combined number (black) during the simulation. The dashed lines correspond to the theoretical mean population sizes in the quasi-stationary state. (a) if the proliferation advantage of the mutant is below the threshold the mutant will rapidly escape from the shallow quasi-stationary state and go extinct. On the bottom panels the black × on the continuous approximation of the potential marks the quasi-stationary state. Parameters are: *N*^0^ = 100, *β* = 1, *r*^+^ = 0, and *S*_m_ = 0.2 is below the threshold 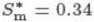. (b) Increasing the compartment size to *N*^0^ = 400 the potential well becomes deeper and the threshold proliferation advantage correspondingly smaller 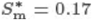, allowing a mutant with the same advantage of *S*_m_ = 0.2 to persist in the tissue during the individual’s lifetime.

Here we are concerned with cell number regulation that can be described by a confining potential with a single minimum, for which Eq. (3) is the parabolic approximation. In this case the presence of size regulation (i.e., *β* > 0) leads to a quasi-stationary state in which the mean number of mutant 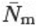 and wild type 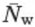 cells can be determined to good approximation by solving:

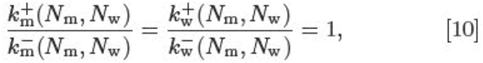

which gives:

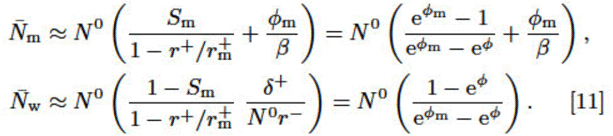

As illustrated in Fig. 2, the behavior of this quasi-stationary state can be divided into two regimes based on the value of the proliferative advantage *S*_*m*_ of the mutant. Below a threshold proliferative advantage 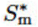 mutants, even if they initially spread (i.e., avoid early stochastic extinction with probability *S*_*m*_) will nonetheless rapidly go extinct and, as a result, have vanishing probability to persist in the tissue throughout its lifetime. Above this threshold, however, if a single mutant avoids early stochastic extinction, with probability *S*_*m*_, a population of its descendants will persist with near certainty in the tissue throughout its lifetime.

The characteristic residence time of a population of mutants that have initially spread corresponds to the mean exit time *τ* of escape from the quasi-stationary state described above. Following the approach described by Gardiner (17) and Derényi et al. (18) an analytical approximation can be derived for *τ* of the general form (for details of see the Supplementary Appendix):

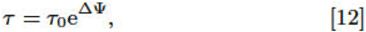

where *τ*_0_ is the reciprocal of the attempt frequency and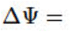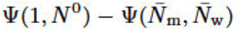 is the height of the potential barrier for the escape from the quasi-stationary state (cf. Fig. 2). **ΔΨ** scales linearly with *N*^0^ (for large *N*^0^), corresponding to an exponential increase in *τ*. In contrast, *τ*_0_, which depends only on the local geometries of the potential well and barrier, is proportional to (*N*^0^)^3/2^

Using the mean exit time for escape from the quasi-stationary sate, the probability ***P*** that a single mutant persists (i.e., first spreads, and then avoids escape) for the lifetime of the individual can be expressed as:

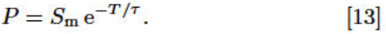

As show in Fig. 3a top panel, the above approximation for the escape time *τ* is highly accurate, and it depends very sharply on the selective advantage of mutants. This results in a well defined threshold selective advantage (**cf.** Fig. 3a bottom panel) below which mutants, even if they avoid early stochastic extinction, will rapidly go extinct, i.e., will be washed out by cells differentiating from below. Furthermore, the threshold value depends only weakly on the value of *β* for reasonably strong cell number regulation, i.e., for *β* > 1 corresponding to the variance (in time) of the cell number being smaller than *N*^0^

**Fig. 3.**
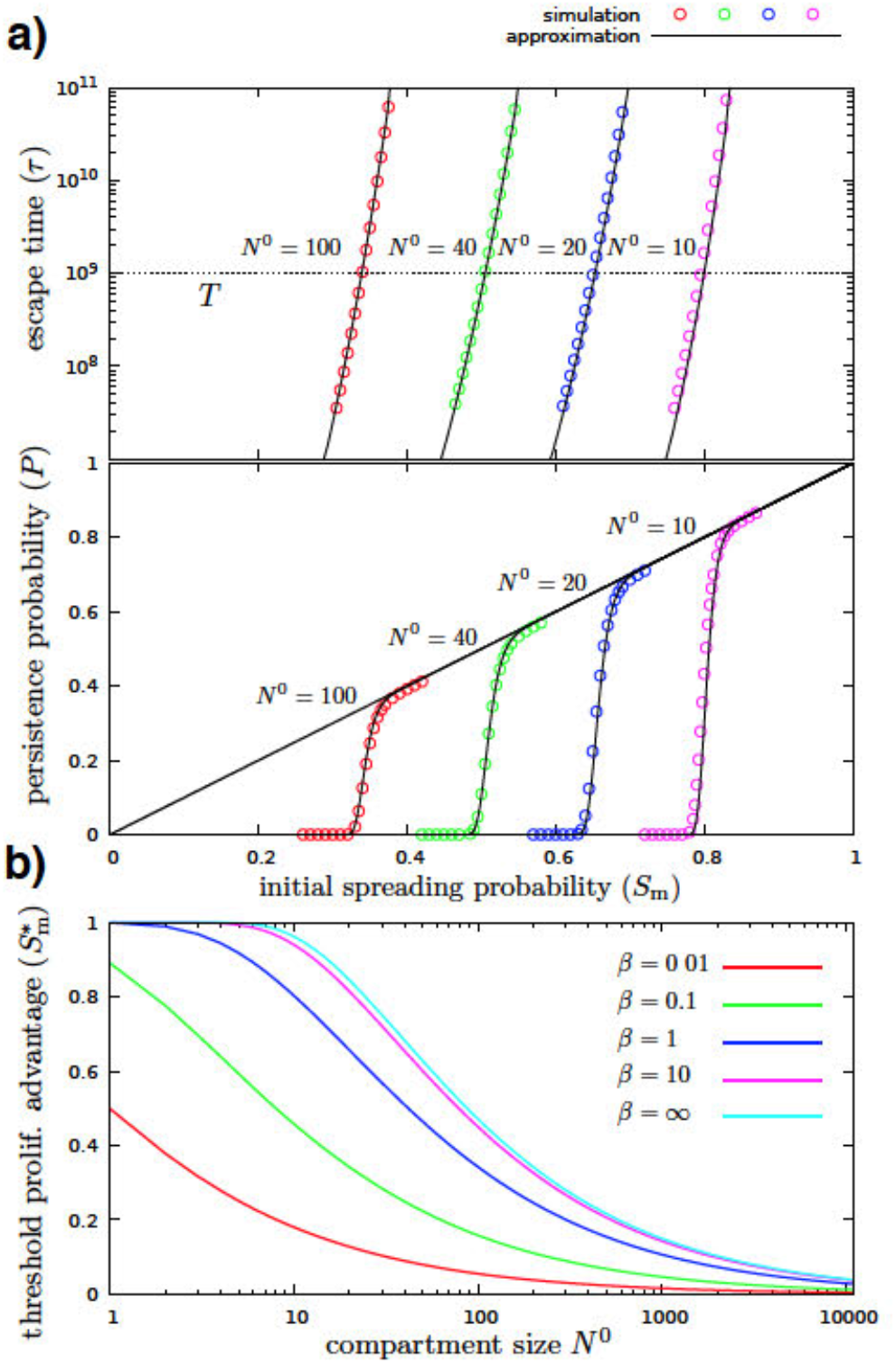
There is no somatic evolution under the threshold spreading factor. (a) In presence of tissue size regulation (*β* > 0), below a threshold proliferation advantage 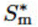 mutants rapidly go extinct and, as a result, have vanishing probability to persist in the tissue through its lifetime, while above this threshold, if a single mutant avoids early stochastic extinction, which occurs with probability *S*_*m*_, it will persist with near certainty. The diagonal line corresponds to the initial spreading probability, the colored circles show the results of the simulation, and the black continuous curves show the theoretical approximation, for different compartment sizes (*N*^0^). *β* = 1, *r*^+^ = 0, and *T* = **10**^9^ throughout (b) The threshold separates the plot into two distinct regimes: below the curve the persistence probability is zero, mutations cannot accumulate; above the curve the mutants that avoid early stochastic extinction, which occurs with probability *S*_*m*_, will persist in the tissue during the lifetime of the individual, and mutations can accumulate, leading to neoplastic progression.

Realistic values for the rates *r*^−^, *r*^+^ and *δ*^+^ can vary greatly depending on tissue type and differentiation state (e.g., in humans long term stem cells of the hematopoietic system divide a few times a year, while in the top layers of epithelial tissues cell divisions occur daily (8, 12)). Under homeostatic conditions the three rates, however, cannot be chosen independently, but must satisfy the stationary condition in Eq. (1). Furthermore, in the context of our model, as is apparent on inspection of _Ψ_, the dynamics does not depend on the absolute rates, but only on the ratio *r*^+^/*r*^−^ the logarithm of which defines the self-renewal potential *ϕ* (cf. Eq. (5)).

Fixing *r*^−^ specifies the absolute time scale, while changing the value of, e.g., *r*^+^ changes the values of the self-renewal potential *ϕ* = ln(*r*^+^/*r*^−^) and the strength of washing out, defined as the fraction of cells being produced by differentiation from below instead of self-renewal: *δ*^+^/(*N*^0^*r*^−^) = 1−*r*^+^/*r*^−^ = 1 - exp(*ϕ*). In particular, *r*^+^ = 0, the default value used in several examples above, corresponds to minimal self-renewal potential (*ϕ* = −∞), and maximal washing out (δ^+^/(*N*°*r*^−^) = 1). Increasing values of *r*^+^/*r*^−^ > 0 correspond to increasing self-renewal potential and weakening washing out. As shown in Fig. S4, even for strong self-renewal and correspondingly weak washing out the threshold spreading factor can be large in small compartments.

## 2. Discussion

In classical population genetics models of finite populations, a mutation is either fixed in the population or lost from it within a finite length of time. A fundamental result of population genetics theory is that in constant populations mutations with a given selective advantage will avoid early stochastic extinction and fix with a probability independent of population size and proportional to the selective advantage (14–16). As a corollary, in the context of somatic evolution Michor et al. (19) demonstrated that the accumulation of oncogene-activating mutations (i.e., mutations that provide a proliferative advantage) that occur at a constant rate per cell division is faster in large than in small compartments. Consequently, as pointed out by Michor et al. the classical theory of finite populations of constant size implies that the organization of self-renewing tissues into many small compartments, such as the stem cell pools in colonic crypts, from which the tissue is derived, protects against cancer initiation (5). Further work by Beerenwinkel et al. using qualitatively similar models with a single compartment without differentiation from below, found that the average waiting time for the appearance of the tumor is strongly affected by the selective advantage, with the average waiting time decreasing roughly inversely proportional to the selective advantage. The mutation rate and the size of the population at risk in contrast were found to contribute only logarithmically to the waiting time and hence have a weaker impact (6).

In hierarchically organized tissues with finite compartment size the situation is more complicated. A mutant that avoids early stochastic extinction and achieves a sizable seemingly stable population can go extinct as a result of differentiation from below. This results in a qualitatively different and more profound ability of smaller compartment size to limit the accumulation of mutations. Similarly to classical population genetics models, the initial spreading probability of a mutation in a compartment of a hierarchical tissue is proportional to the proliferative advantage *S*_m_ and independent of the compartment size. However, as can be seen in Fig. 3a, the probability of the mutation to persist in the tissue exhibits a threshold that is strongly dependent on compartment size. For small compartments even mutants with a very large selective advantage will only persist for a very short time, e.g., a mutant with a selective advantage of 10%, i.e., *S*_m_ ≈ 0.1, the largest value considered by Beerenwinkel et al., will rapidly go extinct in compartments with up to several hundred cells.

An important exception is constituted by tissue specific stem cell compartments residing at the bottom of the hierarchy, such as the stem cells at the bottom of colonic crypts. As these compartments do not receive an influx of cells from lower levels, their dynamics can be described by the classical population genetics models discussed above, i.e., mutations can accumulate more easily.

The derivation of the results presented above relies on the existence of the potential defined in Eq. (9). In our model this is ensured by the assumptions that (i) the transition rates for cells of each type depend only on the number of cells of that type; and (ii) cell number regulation acts as a multiplicative rate modifier and depends only on the total number of cells. There are several biologically relevant violations that must be considered. In real tissues the first assumption, the independence in the absence regulation, is in general violated by mutation of wild type cells into mutant cells (and vice versa), as this increases the number of mutant cells at a rate dependent on the number of wild type cells (and vice versa). In the context of most, if not all, somatic tissues the rate of mutations that confer significant selective advantage is sufficiently low that the waiting time between successive mutations is much longer than the relevant time scale of the dynamics considered here, thus, it has a negligible effects on the persistence time and, as a result, it does not effect our conclusions. This is even more so the case for back mutations from mutants to the wild type. The second assumption, the postulation of a simple form of cell number regulation that acts as a multiplicative modifier and depends only on the total number of cells, is clearly a simplification. It neglects, for instance, explicit spatial organization and any potential long term memory, such as hysteresis of the homeostatic compartment size dependent on either intrinsic or extrinsic parameters. Such a simplified form of regulation, however is consistent with more detailed models of homeostatic tissue size regulation, such as recent work on the stability of regulation (20–22) and its optimality in terms of reducing mutation accumulation (23).

In order to quantitatively discuss the biological relevance of our results we must consider relevant values of two parameters: compartment size (*N*^0^) and the strength of the homeostatic cell-number regulation (*β*). Consider for instance the intestinal crypts. Our knowledge of intestinal crypt organization is most extensive for murine tissues where crypts are believed to consists of approximately 250 cells in total, out of which 160-180 are proliferative progenitor cells and 4-8 are stem cells residing near the bottom of the crypt (24–27). Methods using bromodeoxyuridine labeling (28), Ki-67 antibody staining (20), and analysis of methylation patterns (29) conclude that the crypts in humans contain around 2000 cells with the number of progenitor cells being between 500 and 700. In the context of our model, assuming that proliferative cells can be regarded as belonging to between 1 10 discrete levels of progressively faster dividing cells corresponds to values of *N*^0^ ≈ 170 − 17 cells in mice and *N*^0^ ≈ 600 − 60 in humans. Experimental evidence on the strength of cell number regulation is much more limited. Bravo and Axelrod (20), however have measured the variation in cell numbers across biopsies in 49 crypts from human individuals and found a mean of 624 proliferative cells with a standard deviation of 234. Assuming that (i) all the proliferative cells belong to a single compartment and (ii) all of the observed variation across crypts can be attributed to cell-number fluctuations around a common homeostatic value, i.e., ignoring completely variation in homeostatic size across crypts and neglecting measurement error, provides a lower bound on the strength of regulation of *β* ≪ 624/234^2^ ≈ 0.01. The generally well-defined cylindrically symmetric morphology of crypts, however, suggests that a standard deviation corresponding to at most 10% of the mean cell number is more realistic. Assuming between 1 − 10 levels this corresponds to 1/(62.4 × 0.1^2^) ≈ 1.6 > *β* > 1/(624 × 0.1^2^) ≈ 0.16 in human and 1/(17 × 0.1^2^) ≈ 6 > *β* > 1/(170 × 0.1^2^) ≈ 0.6 in mouse. This, together with the above values for *N*^0^ places the threshold selective advantage at between 0.1 and 0.5 in the human colon and between 0.15 and 0.7 in mouse (cf. Fig. 3b).

At present systematic data on the selection advantage of mutations in somatic tissues is not available. Vermeulen et al. (7), however, measured the fixation probability of several known drivers of colorectal cancer in the mouse intestine, finding values between 0.4 (Kras +/−) and 0.75 (Kras G12D), which are consistent with the above estimates. In the context of a different epithelial tissue, the human esophagus, a survey by Martincorena et al. of clones persisting in normal tissue showed genomic evidence of strong selective advantage of mutations(30), again consistent with our predictions. Future data on tissue organization and the selection advantage of mutations that persist in normal tissue will offer exciting opportunities to confront them with our results.

## 3. Methods

Detailed derivation of the results presented above are provided in the Supplementary Appendix. All data is contained in the manuscript text and Supplementary Appendix.

## ACKNOWLEDGMENTS

GJSz received funding from the European Research Council under the European Union’s Horizon 2020 research and innovation programme under grant agreement no. 714774 and the grant GINOP-2.3.2.-15-2016-00057.

## Supplementary Information

## Supporting Information Text

### Sufficient conditions for conservative cell number dynamics

**Fig. S1.**
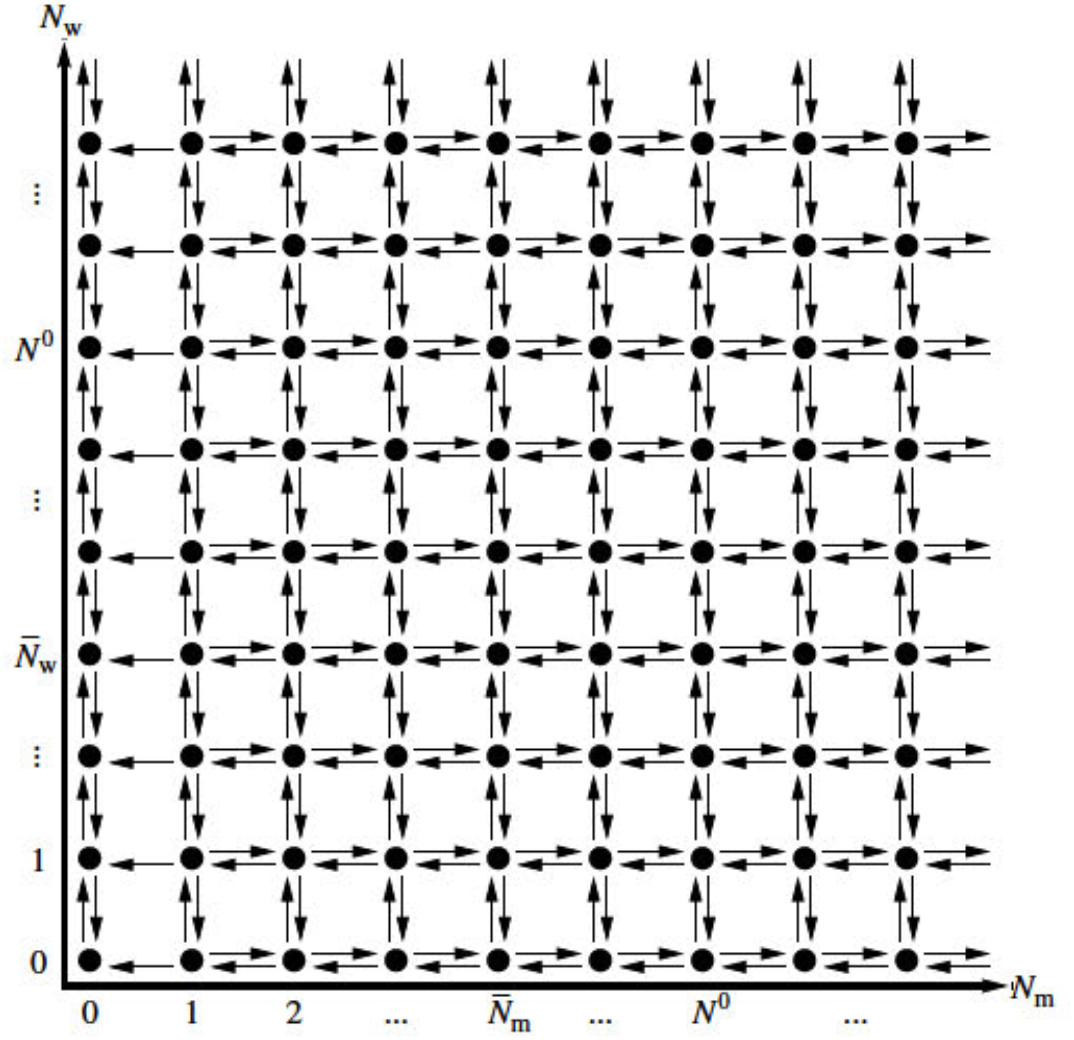
**The state space**, where the allowed states (*N*_m_, *N*_w_) are denoted by black circles, and the transitions with non-zero rates by black arrows.

Consider the cell number dynamics where (i) both the mutant and wild type transition rates depend only on the number of cells of the given type; and (ii) cell number regulation acts as a multiplicative modifier of these rates and depends only on the total number of cells:

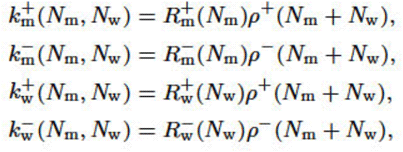

where 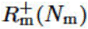 and 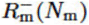 indicate the rates of the transitions in the number of mutant cells: *N*_m_ ➔ *N*_m_ +1 and *N*_m_ ➔ *N*_m_ − 1, respectively, and 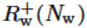 and 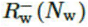 indicate the rates of the transitions of wild type cells: *N*_w_ ➔ *N*_w_ + 1 and *N*_w_ ➔ *N*_w_ − 1. The state space of the tissue compartment under consideration is depicted in Fig. S1, where all the transitions with nonzero rates are indicated by arrows. Starting from an arbitrary population state *N*_m_ = *i*, *N*_w_ = ***j*** the product of the equilibrium constants (i.e., the ratios between to forward and backward rates) along an elementary cycle (*i*,*j*) ➔ (*i*,*j*+1) ➔ (*i*−1,*j*+1)➔ (*i*−1,*j*) ➔ (*i*,*j*) is unity:

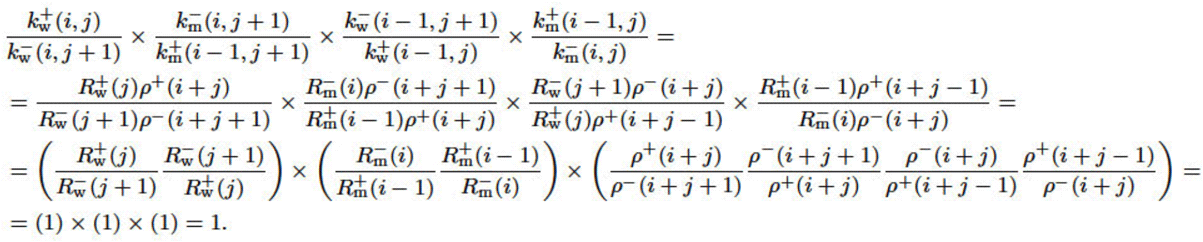

This is equivalent to Kolmogorov’s criterion, and it follows that the cell number dynamics is conservative and described by a potential.

More generally, consider an evolutionary dynamics with an arbitrary number of cell types in a potential. Similarly to the above derivation, it can be shown that if the cell number increasing and decreasing rates for any subset of cell types are multiplied by, respectively, cell number increasing and decreasing rate modifiers that depend only on the total number of cells of the given subset, then the resulting dynamics is also described by a potential. Moreover, the resulting potential is the sum of the two potentials corresponding to the original dynamics and to the rate modifiers. This is because (i) along any path in the state space the product of the equilibrium constants can be factored into the products of the equilibrium constants corresponding to the original dynamics and to the rate modifiers; and (ii) along any elementary cycle both products are unity. As a corollary, if the transition rates of an evolutionary dynamics are factorizable into functions such that each function corresponds to the increase or decrease of the number of cells of a subset of cell types, and each function depends only on that cell number, then the dynamics is conservative and described by a potential.

### Derivation of the potential Ψ(*N*_m_, *N*_w_)

For the transition rates considered in the main text, the potential Ψ(*N*_m_, *N*_w_) corresponding to the dynamics can be calculated as the logarithm of the product of the equilibrium constants from point (*N*_m_, *N*_w_) to a reference point, e.g., to the lower left state (1, 0) of the reversible part of the state space, up to an additive constant. Taking the path (*N*_m_, *N*_w_) → (1, *N*_w_) → (1, 0), this product can be written as:

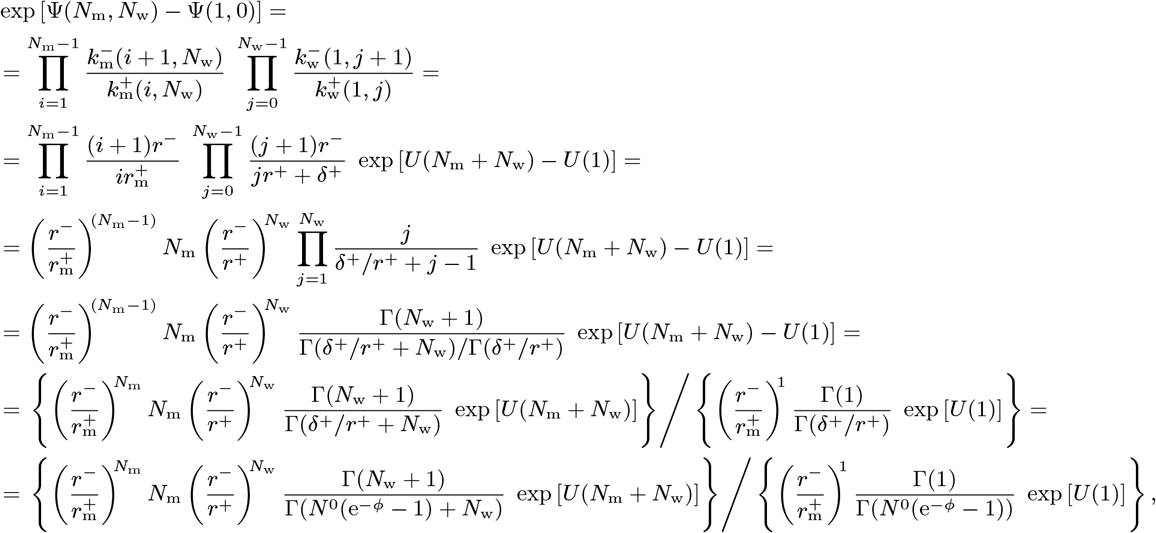

where Γ represents the gamma function and in the last step we use equations (1) and (5) of the main text. The logarithm of the numerator (between the braces) results in the formula for Ψ(*N*_m_, *N*_w_) given in Eq. (9) of the main text, while the logarithm of the denominator is identical to Ψ(1, 0).

### Mean exit time

The derivation of the mean exit time from the effective potential well near the quasi-stationary state 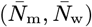 to the boundary line corresponding to zero mutants (*N*_m_ = 0) is outlined below. Exact analytical formula exists either for continuous systems of arbitrary dimensions (subsections 5.2.7 and 9.3.2 of Gardiner (1)), or discrete systems in one dimension (section 7.4 of Gardiner (1) and Derényi et al. (2)). One can, however, generalize the discrete one-dimensional formula to our discrete two-dimensional system in a straightforward manner.

Let us first select any of the shortest paths (with only upward and leftward transitions) form 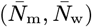 through (1, *N*^0^) to (0, *N*^0^). The main contribution to the mean exit time (which, in one dimension, is often referred to as “mean first passage time”) along this one-dimensional path comes from the product of all backward transition rates (except for the outermost one from (0, *N*^0^) to (1, *N*^0^)) divided by the product of all forward transition rates (including the one from (1, *N*^0^) to (0, *N*^0^)). This contribution is independent of the selected path (because the rates correspond to an effective potential, Ψ(*N*_m_, *N*_w_)):

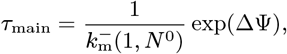

where

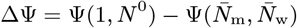

is the height of the effective potential barrier against escape.

The mean exit time involves the sum of similar contributions between any pairs of states along the selected path (section 7.4 of Gardiner (1) and Derényi et al. (2)). Only those contributions are significant for which the starting and ending positions are close to the bottom and the top of the effective potential, respectively. The summation for these contributions leads to a correction factor to the main contribution. This correction factor consists of two terms: a sum of the 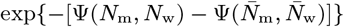 Boltzmann weights of the states (*N*_m_, *N*_w_) along the path near the bottom; and a conceptually similar sum near the top (not detailed here, because its terms cannot be readily expressed by the potential w, but rather only by the products of the ratios of the corresponding transition rates).

Because a typical exit process follows the diagonally oriented potential valley of the state space (see Fig. 2 in the main text), let us restrict the selected path to run along this valley. The correction factor for such a path will depend only on the local “geometry” of the bottom and the top of the effective potential, parallel to the direction of the main valley. In particular, the correction factor at the top (which is the effective width of the potential barrier) can be approximated as

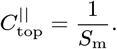

This one-dimensional result can be generalized to two (or any higher) dimensions in analogy to the generalization of the one-dimensional continuous version of the mean exit time (see subsection 9.3.2 of Gardiner (1)): the sum of the above Boltzmann weights should be extended to all states near the bottom of the effective potential:

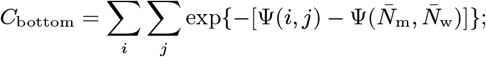

and the result should be divided by the sum of a different type of Boltzmann weights for all the states along the exit line, *N*_m_ =1 (which gives the effective width of the potential saddle at the barrier):

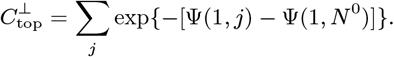

The resulting mean exit time

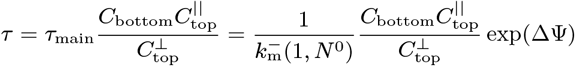

is proportional to exp(ΔΨ) and its prefactor, denoted by *·*0, is often referred to as the reciprocal of the attempt frequency. The summations can be approximated by closed formulas using quadratic approximations for the effective potential near the bottom and the top. However, to achieve higher accuracy we executed the summations numerically for displaying the theoretical estimates in the figures of both the main text and the Supplementary Information.

### Simulations

Two types of simulations were developed to study the cell dynamics and measure the persistence probability of mutants in hierarchically organized tissues:

#### Explicit kinetics

We performed explicit kinetic Monte Carlo simulations (also known as the “Gillespie algorithm”), where the number of mutants *N*_m_ and wild type cells *N*_w_ evolved according to the rates described in Eq. (8) of the main text. At each iteration, rates were calculated based on the current value of *N*_m_ and *N*_w_, one of the four possible events was chosen proportional to its rate, and *N*_m_ or *N*_w_ was changed accordingly, while time was increased by the reciprocal of the sum of the four rates. The simulations were started with *N*_m_ =0 and *N*_w_ = *N*^0^. At time *t* = 20000, measured in units of 1/(2*r*^−^*N*^0^), a single mutant was introduced by setting *N*_m_ = 1. The simulations were stopped after reaching time *T* = 10^9^. Data used in Fig. 2 of the main text were generated using the explicit kinetic simulation.

#### Efficiently measuring the persistence probability

The time evolution of the probability distribution *P*_i,j_(*t*) of the number of (*i* mutant and *j* wild type) cells is described by the master equation defined by the transition rates given in Eq. (8) of the main text. The inward and outward probability currents into and out of state (*i,j*) can be written, respectively, as

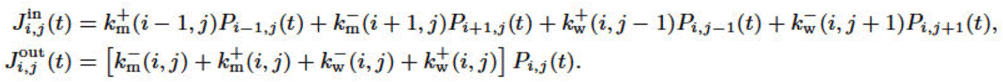

To efficiently measure the spreading probability (*S*_m_) and the mean exit time of escape from the quasi-stationary state (τ), we calculated the steady state probability distributions for the following two modified process:

##### Process I

The process starts with a single mutant and *N*° wild type cells (green circle in Fig. S2), and each time mutants would either go extinct (red arrows in Fig. S2) or spread and reach their quasi-stationary number 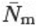 (blue arrows in Fig. S2), the process is restarted in the (1, *N*^0^) state.

**Fig. S2.**
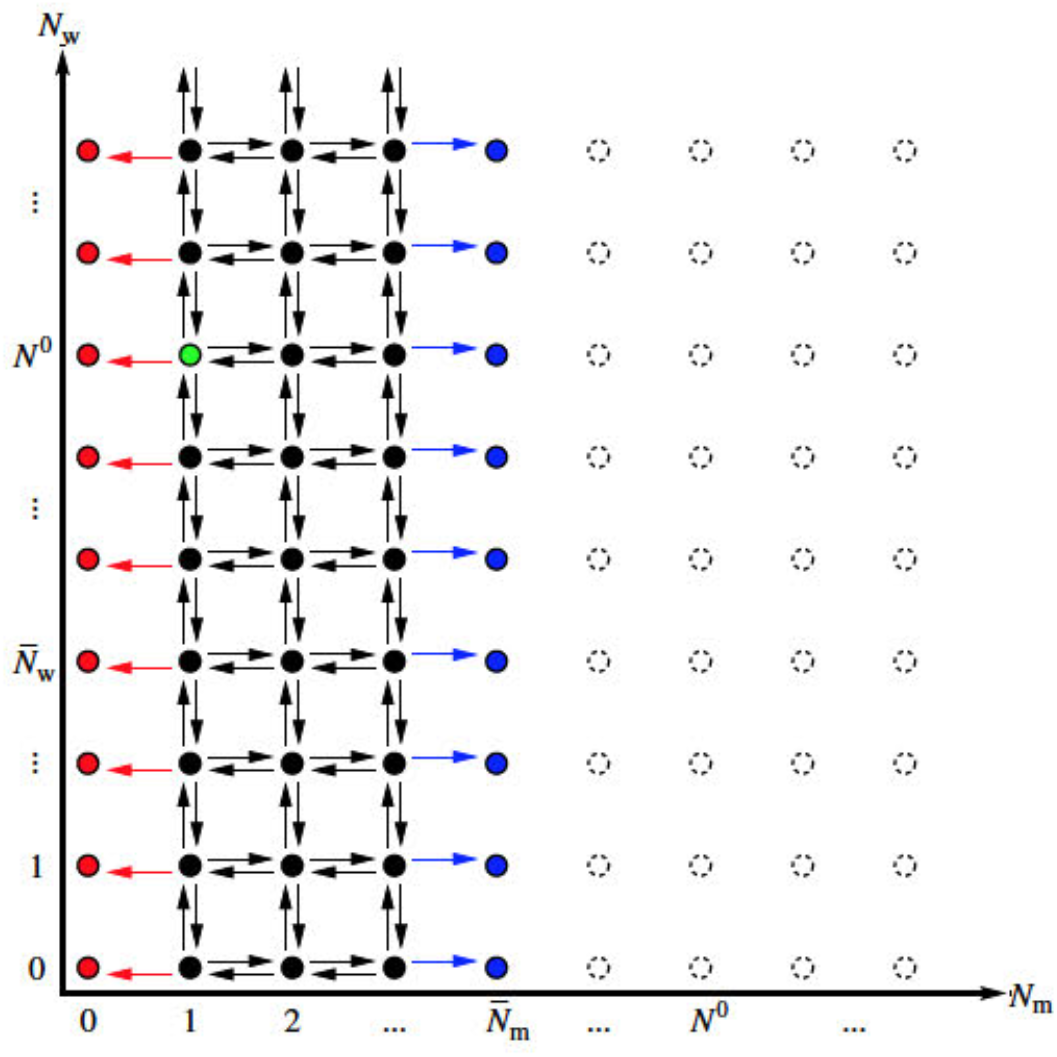
The state space corresponding to Process I.

The extinction and spreading currents can be defined, respectively, as:

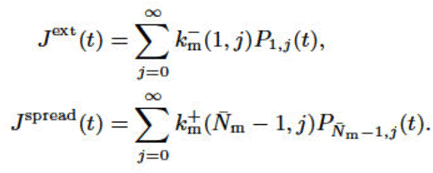

Thus the master equation of the process for the states with more than 0 but less than 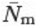 mutants (solid black and green circles in Fig. S2) is:

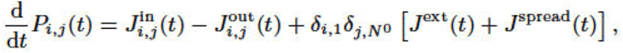

where the Kronecker delta symbol *δ*_j,i_ is 1 if *j* = *i*, and 0 otherwise.

Numerically solving the master equation (using Euler’s method and setting a large enough upper bound for the number of wild type cells), the probability distribution together with the extinction and spreading currents converge to their steady state values, denoted by 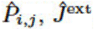 and 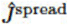, respectively. The probability that a single mutant can spread and avoid early stochastic extinction is, thus:

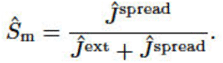

**Fig. S3.**
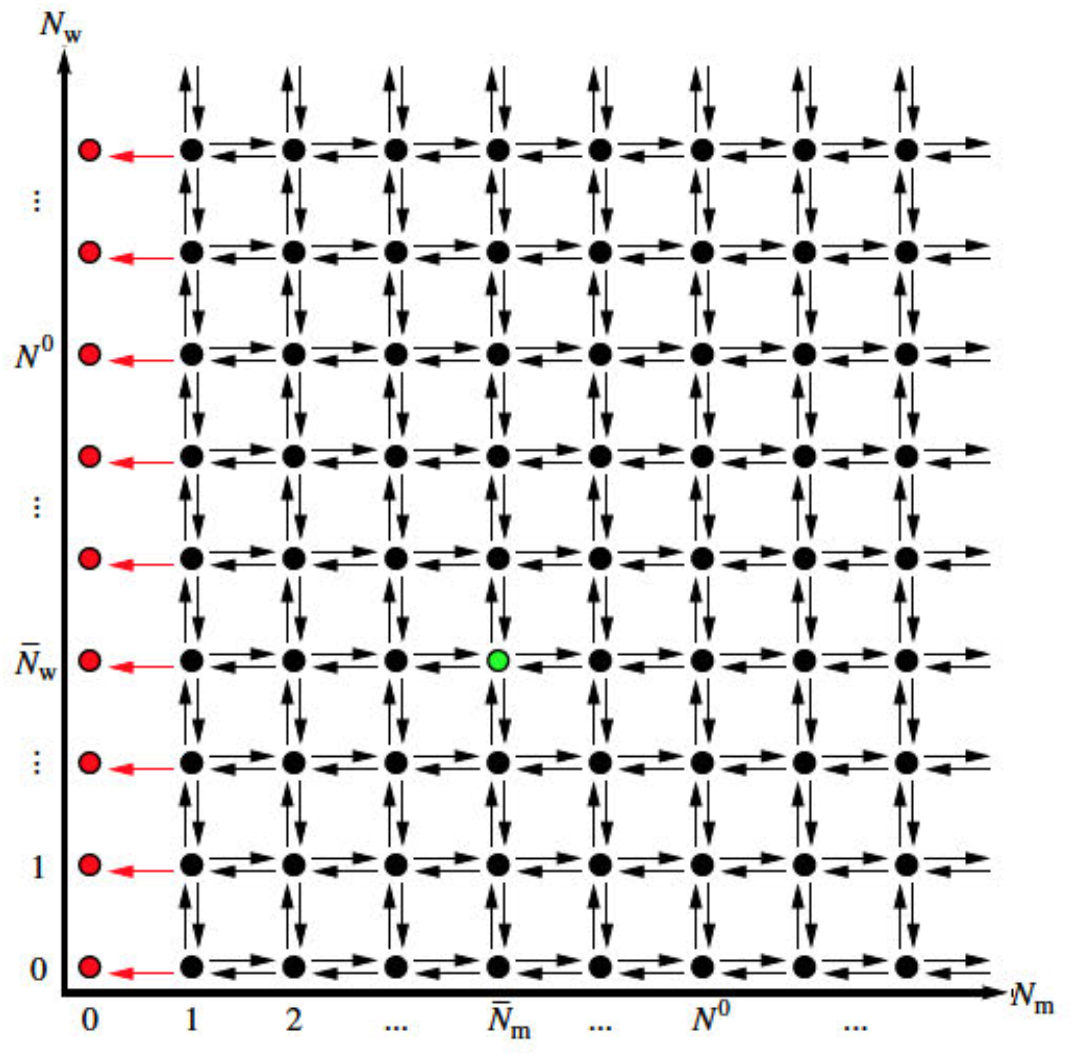
The state space corresponding to Process II.

##### Process II

The process starts in the quasi-stationary state (green circle in Fig. S3), i.e., with 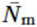 mutants and 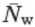 wild type cells, and each time mutants would go extinct (red arrows in Fig. S3) the process is restarted in the quasi-stationary state 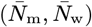,

With the extinction current

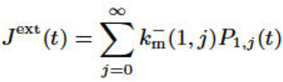

the master equation of the process for all the states with non-zero mutants is:

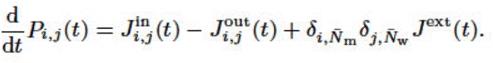

Numerically solving the master equation, the probability distribution together with the extinction current converge to their steady state values 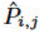 and 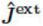, respectively. The reciprocal of the steady state extinction current provides the mean exit time of the mutants from the quasi-stationary state:

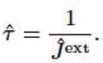

##### Persistence probability

With the combination of the above two processes the probability *P* that a single mutant persists (i.e., first spreads, and then avoids escape) for the lifetime *T* of the individual can be expressed as:

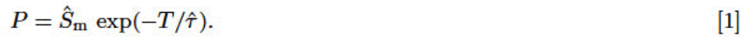

Data used for Fig. 3 of the main text and Fig. S4 below were produced by this method.

**Fig. S4.**
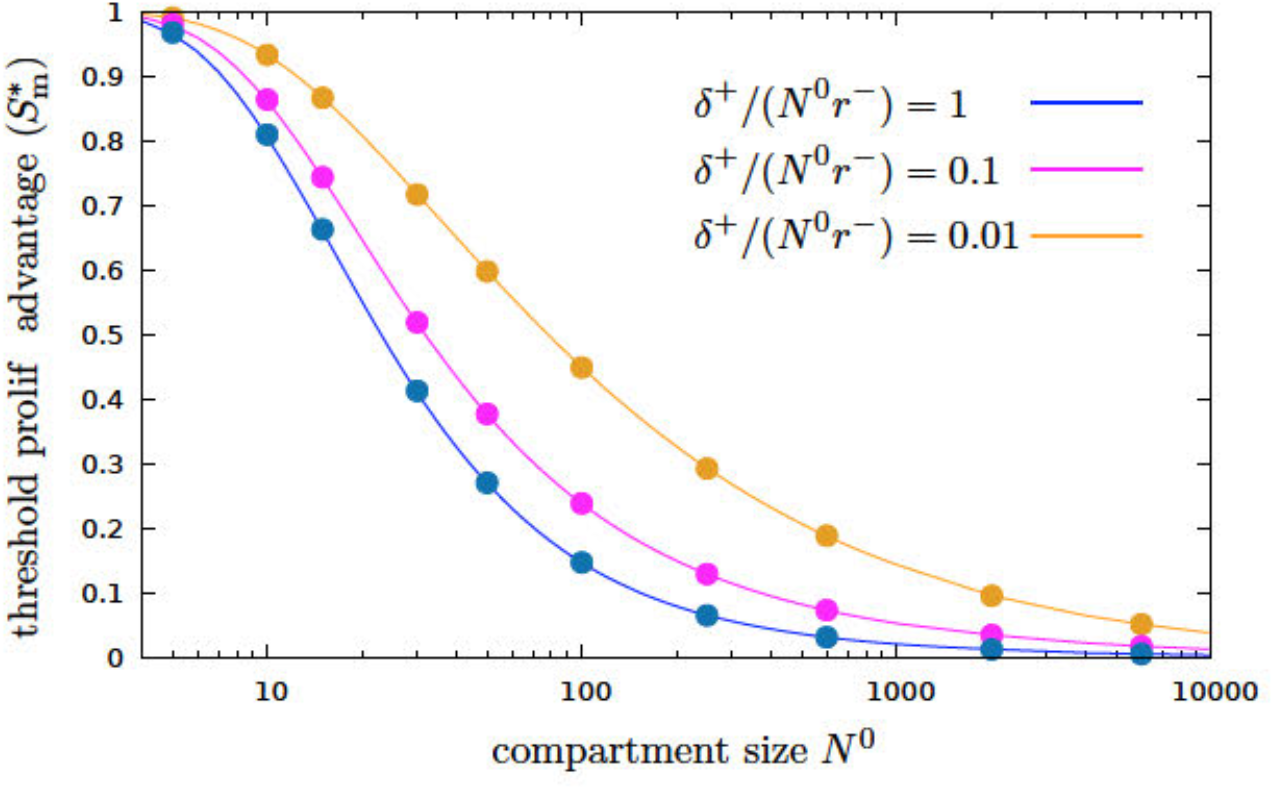
**The threshold spreading factor tor varying strength of washing out**, corresponding to δ^+^/(*N*^0^*r*^−^) = 1, 0.1, and 0.01, i.e., to 100%, 10%, and 1% of cells being produced by differentiation from below instead of self-renewal or, equivalently, *ϕ*, = 1 − ln[δ^+^/(*N*^0^*r*^−^)] ≈ −∞, −0.1, and −0.01, respectively, for *β* = 10 and *T* = 10^9^. Similarly to Fig. 3b in the main lext the threshold spreading factor separates the plot Into two distinct regimes: below the curve the persistence probability is close to zero, mutations cannot accumulate; while above the curve mutants that avoid early stochastic extinction, which occurs with probability Sm, will persist in the tissue during the lifetime of the individual, and can accumulale further mutations leading to neoplastic progression. Continuous lines are theoretical estimates based on the mean exit time approximation, while points indicate explicit numerical simulations.

## References

1. Peter C Nowell. The clonal evolution of tumor cell populations. Science, 194(4260):23–28, 1976.

2. Philipp M Altrock, Lin L Liu, and Franziska Michor. The mathematics of cancer: integrating quantitative models. Nature Reviews Cancer, 15(12):730, 2015.

3. Ivana Bozic, Tibor Antal, Hisashi Ohtsuki, Hannah Carter, Dewey Kim, Sining Chen, Rachel Karchin, Kenneth W Kinzler, Bert Vogelstein, and Martin A Nowak. Accumulation of driver and passenger mutations during tumor progression. Proceedings of the National Academy of Sciences, 107(43):18545–18550, 2010.

4. Christopher D McFarland, Leonid A Mirny, and Kirill S Korolev. Tug-of-war between driver and passenger mutations in cancer and other adaptive processes. Proceedings of the National Academy of Sciences, 111(42):15138–15143, 2014.

5. Niko Beerenwinkel, Roland F Schwarz, Moritz Gerstung, and Florian Markowetz. Cancer evolution: mathematical models and computational inference. Systematic biology, 64(1):e1–e25, 2014.

6. Niko Beerenwinkel, Tibor Antal, David Dingli, Arne Traulsen, Kenneth W Kinzler, Victor E Velculescu, Bert Vogelstein, and Martin A Nowak. Genetic progression and the waiting time to cancer. PLoS computational biology, 3(11):e225, 2007.

7. Louis Vermeulen, Edward Morrissey, Maartje Van Der Heijden, Anna M Nicholson, Andrea Sottoriva, Simon Buczacki, Richard Kemp, Simon Tavaré, and Douglas J Winton. Defining stem cell dynamics in models of intestinal tumor initiation. Science, 342(6161):995–998, 2013.

8. Cristian Tomasetti and Bert Vogelstein. Variation in cancer risk among tissues can be explained by the number of stem cell divisions. Science (New York, NY), 347(6217):78, 2015.

9. Song Wu, Scott Powers, Wei Zhu, and Yusuf A Hannun. Substantial contribution of extrinsic risk factors to cancer development. Nature, 529(7584):43, 2016.

10. Benjamin Werner, David Dingli, and Arne Traulsen. A deterministic model for the occurrence and dynamics of multiple mutations in hierarchically organized tissues. Journal of The Royal Society Interface, 10(85):20130349, 2013.

11. Martin A Nowak, Franziska Michor, and Yoh Iwasa. The linear process of somatic evolution. Proceedings of the national academy of sciences, 100(25):14966–14969, 2003.

12. Imre Derényi and Gergely J Szölloősi. Hierarchical tissue organization as a general mechanism to limit the accumulation of somatic mutations. Nature communications, 8:14545, 2017.

13. NTJ Biley. The elements of stochastic processes. Wiley. New York, 1964.

14. John Burdon Sanderson Haldane. A mathematical theory of natural and artificial selection, part v: selection and mutation. Mathematical Proceedings of the Cambridge Philosophical Society, 23(7):838–844, 1927.

15. Motoo Kimura and Tomoko Ohta. The average number of generations until fixation of a mutant gene in a finite population. Genetics, 61(3):763, 1969.

16. Zaheerabbas Patwa and Lindi M Wahl. The fixation probability of beneficial mutations. Journal of The Royal Society Interface, 5(28):1279–1289, 2008.

17. C Gardiner. Handbook of stochastic methods for physics, chemistry and natural sciences. Springer. Berlin, 2004.

18. Imre Derényi, Denis Bartolo, and Armand Ajdari. Effects of intermediate bound states in dynamic force spectroscopy. Biophysical journal, 86(3):1263–1269, 2004.

19. Franziska Michor, Yoh Iwasa, and Martin A Nowak. Dynamics of cancer progression. Nature reviews cancer, 4(3):197, 2004.

20. Rafael Bravo and David E Axelrod. A calibrated agent-based computer model of stochastic cell dynamics in normal human colon crypts useful for in silico experiments. Theoretical Biology and Medical Modelling, 10(1):66, 2013.

21. Natalia L Komarova and P Van Den Driessche. Stability of control networks in autonomous homeostatic regulation of stem cell lineages. Bulletin of mathematical biology, 80(5):1345–1365, 2018.

22. Jienian Yang, David E Axelrod, and Natalia L Komarova. Determining the control networks regulating stem cell lineages in colonic crypts. Journal of theoretical biology, 429:190–203, 2017.

23. Cesar Alvarado, Nicole A Fider, Helen J Wearing, and Natalia L Komarova. Optimizing homeostatic cell renewal in hierarchical tissues. PLoS computational biology, 14(2):e1005967, 2018.

24. Christopher S Potten and Markus Loeffler. Stem cells: attributes, cycles, spirals, pitfalls and uncertainties. lessons for and from the crypt. Development, 110(4):1001–1020, 1990.

25. Carmen Pin, Alastair JM Watson, and Simon R Carding. Modelling the spatio-temporal cell dynamics reveals novel insights on cell differentiation and proliferation in the small intestinal crypt. PloS one, 7(5):e37115, 2012.

26. Peter Buske, Jörg Galle, Nick Barker, Gabriela Aust, Hans Clevers, and Markus Loeffler. A comprehensive model of the spatio-temporal stem cell and tissue organisation in the intestinal crypt. PLoS computational biology, 7(1):e1001045, 2011.

27. Nick Barker, Johan H Van Es, Jeroen Kuipers, Pekka Kujala, Maaike Van Den Born, Miranda Cozijnsen, Andrea Haegebarth, Jeroen Korving, Harry Begthel, Peter J Peters, et al. Identification of stem cells in small intestine and colon by marker gene lgr5. Nature, 449(7165): 1003, 2007.

28. Christopher S Potten, M Kellett, Stephen A Roberts, DA Rew, and GD Wilson. Measurement of in vivo proliferation in human colorectal mucosa using bromodeoxyuridine. Gut, 33(1): 71–78, 1992.

29. Pierre Nicolas, Kyoung-Mee Kim, Darryl Shibata, and Simon Tavaré. The stem cell population of the human colon crypt: analysis via methylation patterns. PLoS computational biology, 3 (3):e28, 2007.

30. Iñigo Martincorena, Joanna C Fowler, Agnieszka Wabik, Andrew RJ Lawson, Federico Abascal, Michael WJ Hall, Alex Cagan, Kasumi Murai, Krishnaa Mahbubani, Michael R Stratton, et al. Somatic mutant clones colonize the human esophagus with age. Science, 362(6417): 911–917, 2018.

## References

1. C Gardiner. Handbook of stochastic methods for physics, chemistry and natural sciences. Springer. Berlin, 2004.

2. lmre Derényi, Denis Bartolo, and Armand Ajdari. Effects of intermediate bound states in dynamic force spectroscopy. Biophysical journal, 86(3):1263–1269, 2004.

3. Christopher S Potten and Markus Loeffler. Stem cells: attributes, cycles, spirals, pitfalls and uncertainties. lessons for and from the crypt. Development, 110(4):1001–1020, 1990.

4. Carmen Pin, Alastair JM Watson, and Simon R Carding. Modelling the spatio-temporal cell dynamics reveals novel insights on cell differentiation and proliferation in the small intestinal crypt. PloS one, 7(5):e37115, 2012.

5. Peter Buske, Jörg Galle, Nick Barker, Gabriela Aust, Hans Clevers, and Markus Loeffler. A comprehensive model of the spatio-temporal stem cell and tissue organisation in the intestinal crypt. PLoS computational biology, 7(1):e1001045, 2011.

6. Nick Barker, Johan H Van Es, Jeroen Kuipers, Pekka Kujala, Maaike Van Den Born, Miranda Cozijnsen, Andrea Haegebarth, Jeroen Korving, Harry Begthel, Peter J Peters, et al. Identification of stem cells in small intestine and colon by marker gene lgr5. Nature, 449(7165):1003, 2007.

7. Christopher S Potten, M Kellett, Stephen A Roberts, DA Rew, and GD Wilson. Measurement of in vivo proliferation in human colorectal mucosa using bromodeoxyuridine. Gut, 33(1):71–78, 1992.

8. Rafael Bravo and David E Axelrod. A calibrated agent-based computer model of stochastic cell dynamics in normal human colon crypts useful for in silico experiments. Theoretical Biology and Medical Modelling, 10(1):66, 2013.

9. Pierre Nicolas, Kyoung-Mee Kim, Darryl Shibata, and Simon Tavare. The stem cell population of the human colon crypt: analysis via methylation patterns. PLoS computational biology, 3(3):e28, 2007.

